# The distinct biochemical property enables thymidylate kinase as a drug target and participates in pyrimidine drug sensitivity in *Candida albicans*: Specific features of TMPK in *C. albicans*

**DOI:** 10.1101/466938

**Authors:** Chang-Yu Huang, Yee-Chun Chen, Betty A. Wu-Hsieh, Jim-Min Fang, Zee-Fen Chang

## Abstract

The ability to overcome drug resistance in outbreaks of *Candida albicans* infection is an unmet need in health management. Here, we investigated *CDC8*, which encodes thymidylate kinase (TMPK), as a potential drug target for the treatment of *C. albicans* infection. In this study, we found that the specific region spanning amino acids 106-123, namely, the Ca-loop of *C. albicans* TMPK (CaTMPK) contributes to the hyperactivity of this enzyme compared to the human enzyme (hTMPK) and to the utilization of deoxyuridine monophosphate (dUMP)/ deoxy-5-Fluorouridine monophosphate (5-FdUMP) as a substrate. Notably, CaTMPK but not hTMPK enables dUTP/5-FdUTP-mediated DNA toxicity in yeast. CRISPR-mediated deletion of this Ca-loop in *C. albicans* demonstrated the critical role of this Ca-loop in fungal growth and susceptibility to 5-Fluorouridine (5-FUrd). Moreover, pathogenic and drug-resistant *C. albicans* clones were similarly sensitive to 5-FUrd. Thus, this study not only identified a target site for the development of CaTMPK-selective drugs but also revealed 5-FUrd to be a potential drug for the treatment of *C. albicans* infection.

**Author summary:** The emergence of drug-resistant *C. albicans* strains is a serious medical concern that may be addressed by targeting an essential fungal enzyme. *CDC8* encodes thymidylate kinase (TMPK), which is the key enzyme required for dTTP synthesis and is an essential gene for yeast growth. Therefore, the differences of TMPK between human and *C. albicans* can be a potential drug targeting site. This study defines a specific Ca-loop unique to CaTMPK from *C. albicans*, contributing to hyper-activity over human enzyme (hTMPK). CRSPR-edited deletion of this loop also suppressed the growth of *C. albicans*. Moreover, we present evidence that this loop enables dUMP utilization by CaTMPK, but not hTMPK. CaTMPK is also capable of using 5-FdUMP as a substrate, which contributes to 5-FUrd-mediated toxicity. Importantly, we found that many drug resistant pathogenic *C. albicans* isolates from patients are sensitive to 5-FUrd, which has not been used as a drug against fungal infection.

## Introduction

*Candida albicans* is a yeast species and is the most prevalent fungal pathogen in humans [1]. In general, *C. albicans* growing as budding yeast are tolerated by the host immune system. This fungus becomes pathogenic upon switching to hyphal growth in response to changes in environmental temperature and pH and nutrient limitation [2]. Four classes of antifungal drugs have been developed to treat candidiasis: polyenes, azoles, 5-flucytosine (5-FC) and echinocandins. However, major challenges in candidiasis treatment are the acquisition of drug resistance. It has been suggested that drugs such as azoles generate stress in fungus to drive genome evolution, thus selecting gene mutation for developing drug resistance [3-6]. New therapeutic strategies are needed to overcome these obstacles for clinical treatment during infection outbreaks.

Biochemical differences between hosts and pathogens can be exploited to develop selective drugs that have cytotoxic effects on the pathogen but not the host. We targeted an essential enzyme in *C. albicans*, thereby selectively suppressing fungal growth and overcoming drug resistance. Thymidylate kinase (TMPK) is the key enzyme in the biosynthesis of dTTP, catalyzing the conversion of thymidine monophosphate (dTMP) to thymidine diphosphate (dTDP), which is subsequently phosphorylated by nucleoside diphosphate kinase (NDPK) to form thymidine triphosphate (dTTP) [7, 8]. *CDC8*, which encodes TMPK, is an essential gene in yeast; budding yeasts carrying temperature-sensitive mutations in *cdc8* are not viable at nonpermissive temperatures [9-11]. In *C. albicans*, TMPK (CaTMPK) is also encoded by *CDC8*, but the essentiality of this gene has not been previously characterized. The main catalytic modules of TMPK include the LID region, P-loop, DR motif and elements involved in dTMP binding. The sequences of the P-loop, DR motif and TMP-binding elements are highly conserved among TMPK orthologs, while the sequences of the LID region that contribute to the closed conformation for catalysis are divergent. Given the key function of TMPK in dTTP synthesis and the sequence divergence between humans and pathogens, this study investigated the potential application of CaTMPK from *C. albicans* in antifungal drug development.

It has been shown that CaTMPK in complex with ADP and dTMP (PDB ID: 5UIV) at a resolution of 2.45 Å has a unique surface-exposed loop (termed Ca-loop) [12]. However, the functional significance of the Ca-loop in CaTMPK and the molecular mechanism by which the Ca-loop affects catalysis remained unexplored. Here, our data suggest that the Ca-loop mediates hyperactivity of CaTMPK. CRISPR-mediated deletion of this Ca-loop markedly slowed the growth of *C. albicans*, highlighting this region as a new target site for the design of fungi-specific inhibitors. In addition, we provide in vitro and in vivo evidence that CaTMPK is highly efficient at using deoxy-uridine monophosphate (dUMP) as substrates, provoking a possible role of dUTP in stress-induced genetic instability of fungi. Moreover, CaTMPK is also capable of utilizing deoxy-5-fluorouridine monophosphate (5-FdUMP) as the substrate, thus mediating 5-FUrd toxicity. Finally, we showed that 5-FUrd is useful for the treatment of infections caused by *C. albicans* strains that are resistant to 5-FC and azoles.

## Results

### The biochemical differences between hTMPK and CaTMPK

Members of the TMPK enzyme family have been categorized into type I and type II enzymes. Both hTMPK and CaTMPK are type I enzymes [7]. The sequences in P-loop, TMP binding and DR sites in catalysis are highly conserved between CaTMPK and hTMPK (Fig 1A). The structure-based alignment (PDB: 5UIV and PDB:1E2D) [12, 13] shows that the *Candida*-specific Ca-loop starts from Phe107 to Lys108. We have previously developed a series of hTMPK inhibitors [14, 15]. Here, we showed that two compounds, YMU1 and 3b, at 1 μM effectively suppressed hTMPK activity while at 10μM having no inhibitory activity to CaTMPK (Fig 1B). We compared the enzyme activities of CaTMPK and hTMPK. The kinetic parameters of CaTMPK and hTMPK are shown in Table 1. The k_cat_ of CaTMPK for dTMP was 15-fold higher than that of hTMPK, with similar *K_m_* values observed for ATP and dTMP. Thus, the catalytic efficiency of CaTMPK is much higher than that of hTMPK. Taken together, these results suggest that these two TMPKs exhibit distinct properties.

**Table 1.** Kinetic parameters of hTMPK and CaTMPK in dTMP-based reaction.

| | $K_m^{ATP}$ (μM) | $K_m^{dTMP}$ (μM) | $k_{cat}$ ($\text{min}^{-1}$) | $k_{cat}/K_m^{dTMP}$ ($\mu\text{M}^{-1} \text{min}^{-1}$) |
| --- | --- | --- | --- | --- |
| hTMPK | 25.3 ± 3.2 | 16.3 ± 2.3 | 21.1±0.6 | 1.3±0.2 |
| CaTMPK | 44.3±4.2 | 24.1±3.2 | 320.8±7.1 | 13.5±1.0 |

Reactions were performed as described in Materials and Methods. Different concentrations of dTMP (7.5-500 μM) in the presence of 500 μM ATP or different concentrations of ATP (7.5-500 μM) in the presence of 500 μM dTMP were used to determine $K_m$ and $k_{cat}$ using Michaelis-Menton equation: $v = V_{max}[S]/(K_m + [S])$ . Values are presented as mean ± S.D. from at least three independent experiments.

**Fig 1.**
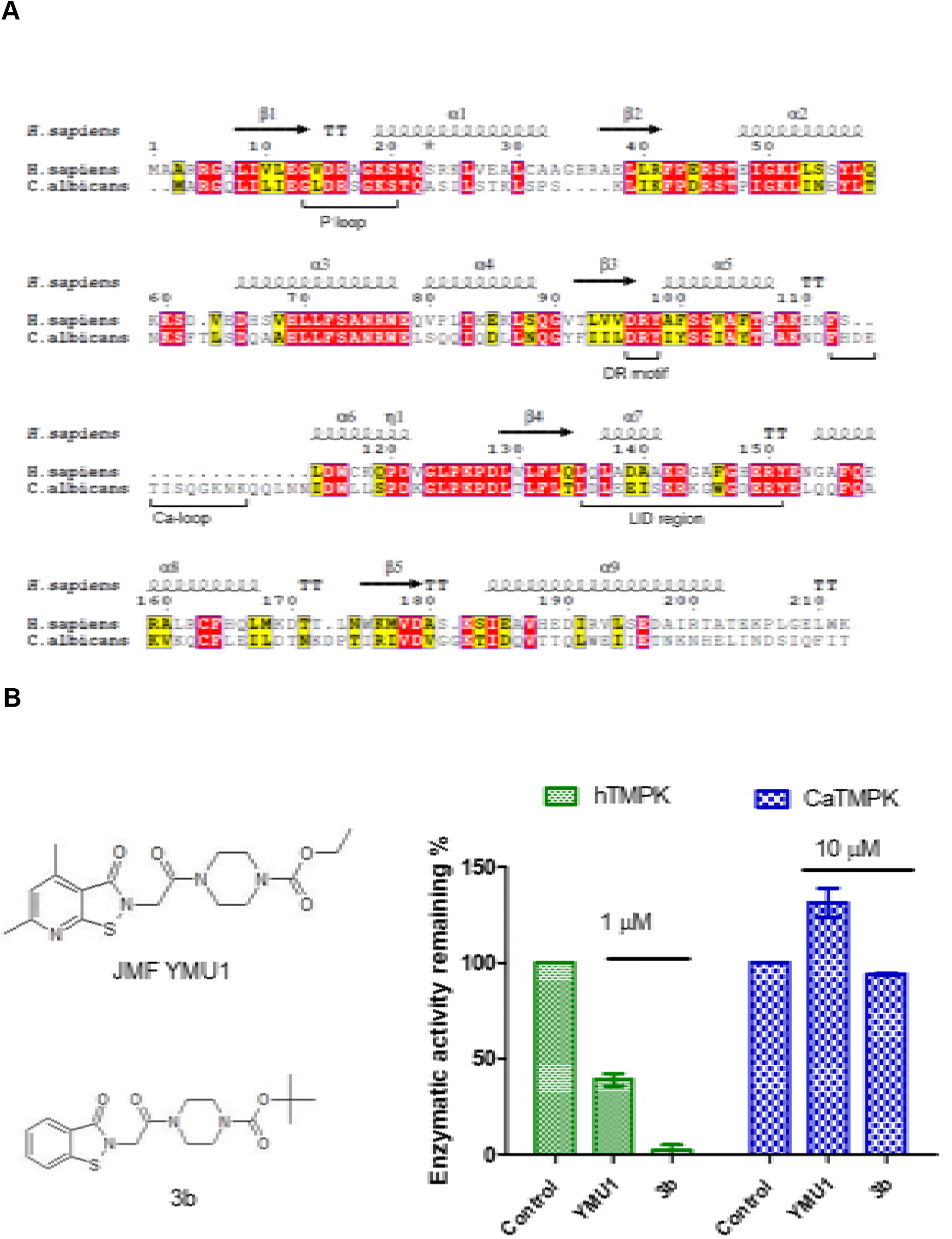
Differences between CaTMPK and hTMPK. (A) Sequence alignment of CaTMPK and hTMPK. Identical and similar residues are shown in red and yellow, respectively. Key catalytic elements, including the P-loop, DR motif and LID region, are labeled. The sequence alignment was generated by MUSCLE [33] and presented by using the ESPript 3.0 program [34]. (B) hTMPK (0.4 μg) and CaTMPK (0.015 μg) was preincubated with hTMPK inhibitors, YMU1 and 3b compounds, at the indicated concentration for 10 min, followed by NADH-coupled assay. Data are expressed in % of activity relative to the control reaction without inhibitor treatment, n = 3. Bar denotes mean ± S.D.

### The Ca-loop confers hyperactivity on CaTMPK and determines the growth rate of *C. albicans*

To verify the contribution of the Ca-loop to the catalytic rate of CaTMPK, we generated a Ca-loop-deleted mutant, Δ107-118. Recombinant wild-type and Δ107-118 CaTMPK proteins were purified for activity assays. The 107-118 deletion led to a 70% reduction in activity (Fig 2A). Because CaTMPK lacking the Ca-loop exhibited decreased activity, we asked whether deletion of this Ca-loop would affect the growth of *C. albicans*. CaTMPK is the gene product of *CDC8*. Therefore, we performed CRISPR-mediated deletion of the 319-354 region of the *CDC8* gene locus (*cdc8Δ107-118*) in the nonpathogenic *C. albicans HLC54* strain [16]. The sgRNA targeting the Ca-loop sequence for Cas9 cleavage and the plasmid pGEX-2T-CaTMPKΔ107-118 as the donor template were cotransformed, followed by selection with nourseothricin (Nat) (S1A and S1B Figs). The selected clone was analyzed by PCR amplification of the region spanning the Ca-loop sequence of the *CDC8* gene locus (S1C Fig.). Sanger sequencing further confirmed that the *cdc8_Δ107-118_* clone had a 36-bp in-frame deletion in the Ca-loop (S1D Fig.). We then compared the growth rates of *HLC54* and *HLC54[cdc8_Δ107-118_]*. The results showed that *HLC54[cdc8_Δ107-118_]* grew much slower than the parental strain *HLC54* (Fig 2B). Taken together, these results show that the Ca-loop is a critical structural element of CaTMPK in the rate of dTDP formation, which is indeed controlling the growth of *C. albicans*.

**Fig 2.**
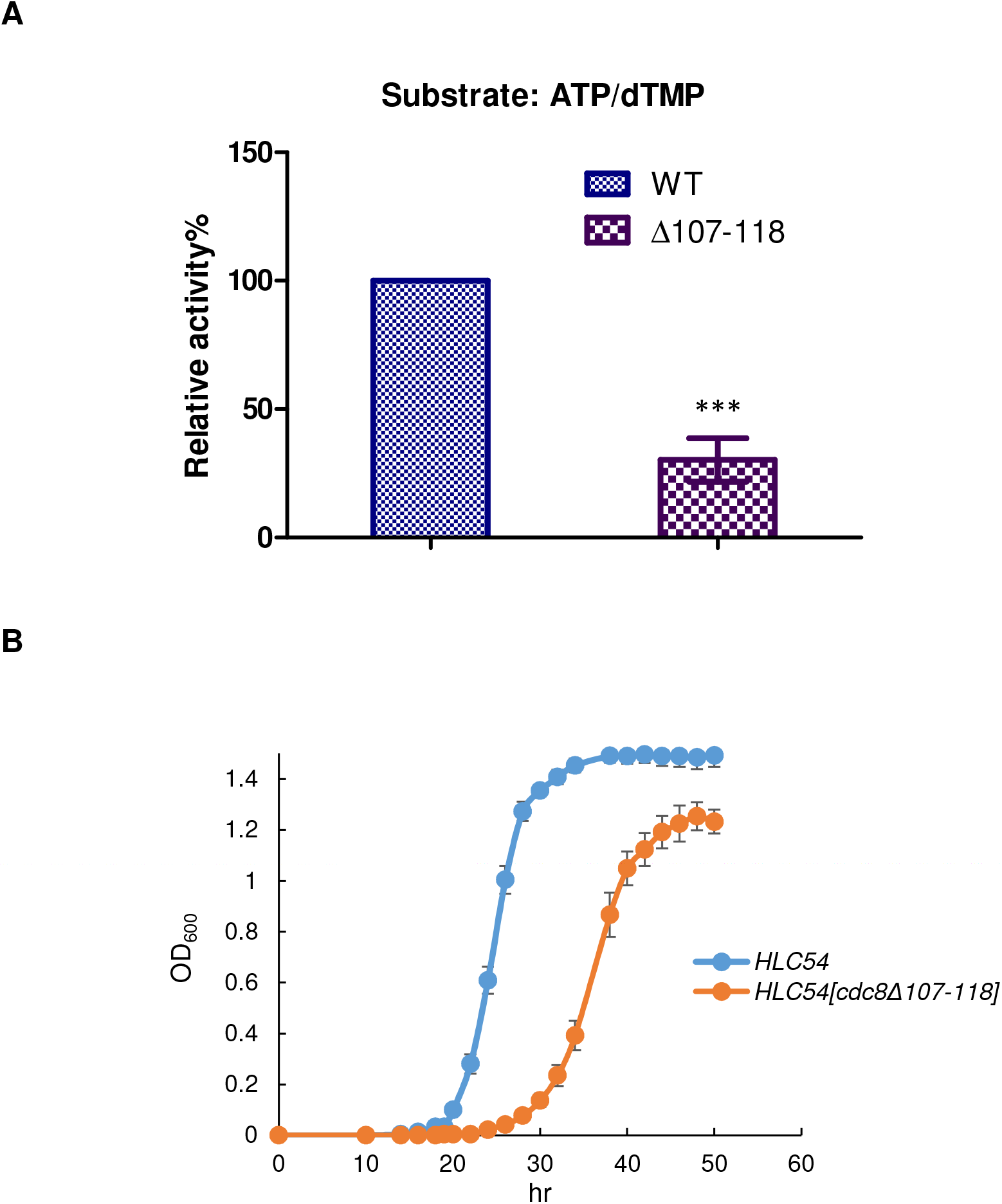
*In vitro* and *in vivo* contribution of the Ca-loop of CaTMPK. (A) TMPK activity of CaTMPK variants. Enzymatic activities of WT and Ca-loop mutants, as indicated, were measured by an NADH-coupled assay in the presence of 500 μM ATP and dTMP. The results are expressed relative to WT CaTMPK activity, which was set to 100%. The results represent the mean ± SD, n=3. P-values were determined by Student’s t-test (*: P<0.05, ***: P<0.001). (B) Effect of CRISPR-based editing of *CDC8* to delete amino acids 107 to 118 on the growth of *C. albicans*. *HLC54* and *HLC54[cdc8*_△*107-118*_] at 10^3^ cells/well in 96-well microplates were incubated at 30°C, and cell densities were determined by measuring the OD_600_ every 30 minutes. Growth curves were constructed for four replicates.

### The Ca-loop allows CaTMPK to use dUMP as a substrate

Given the unique structural feature of the dTMP/LID/Ca-loop of CaTMPK, we further asked whether there is a difference in substrate selectivity between hTMPK and CaTMPK. Here, we performed an isotope-labeling-based γ-phosphate transfer assay to analyze the capacity of purified CaTMPK and hTMPK to phosphorylate different dNMPs. As expected, both enzymes were capable of phosphorylating dTMP to dTDP, and dAMP, dGMP and dCMP were not used as substrates by these enzymes (Fig 3A). However, there was a striking difference in the capability of CaTMPK to transfer γ-^32^P-phosphate from ATP to dUMPs, so as 5-FdUMP (Fig 3B). Analysis of the steady-state kinetics revealed a large difference in the rate of dUMP phosphorylation between hTMPK and CaTMPK (Fig 3C). The k_cat_ of CaTMPK for dUMP was 6-fold higher than that for dTMP, but the catalytic efficiency (k_cat_/*K_m_*) of CaTMPK for dTMP remained higher than that for dUMP (Table 1 and Fig 3C). This finding indicates that CaTMPK normally prefers dTMP as a substrate. However, when the dUMP/dTMP ratio increases, CaTMPK exhibits increased dUDP production. Next, we asked whether the Ca-loop is involved in the utilization of dUMP as a substrate. Similar to the results of the dTMP-based assay, the Δ107-118 was defective in dUMP utilization (Fig 3D), indicating that the Ca-loop determines the utilization of dUMP as a substrate in the enzymatic reaction.

**Fig 3.**
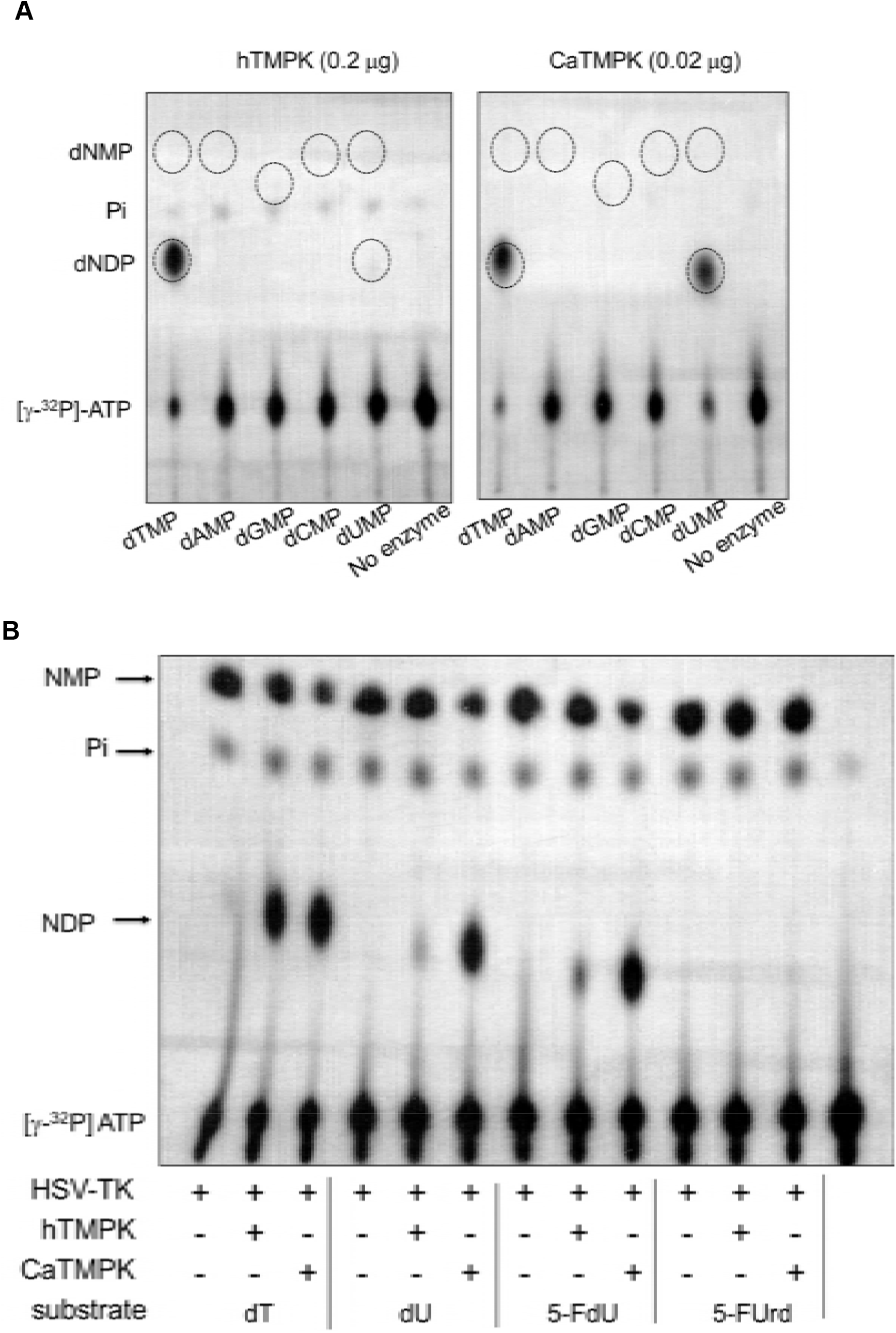
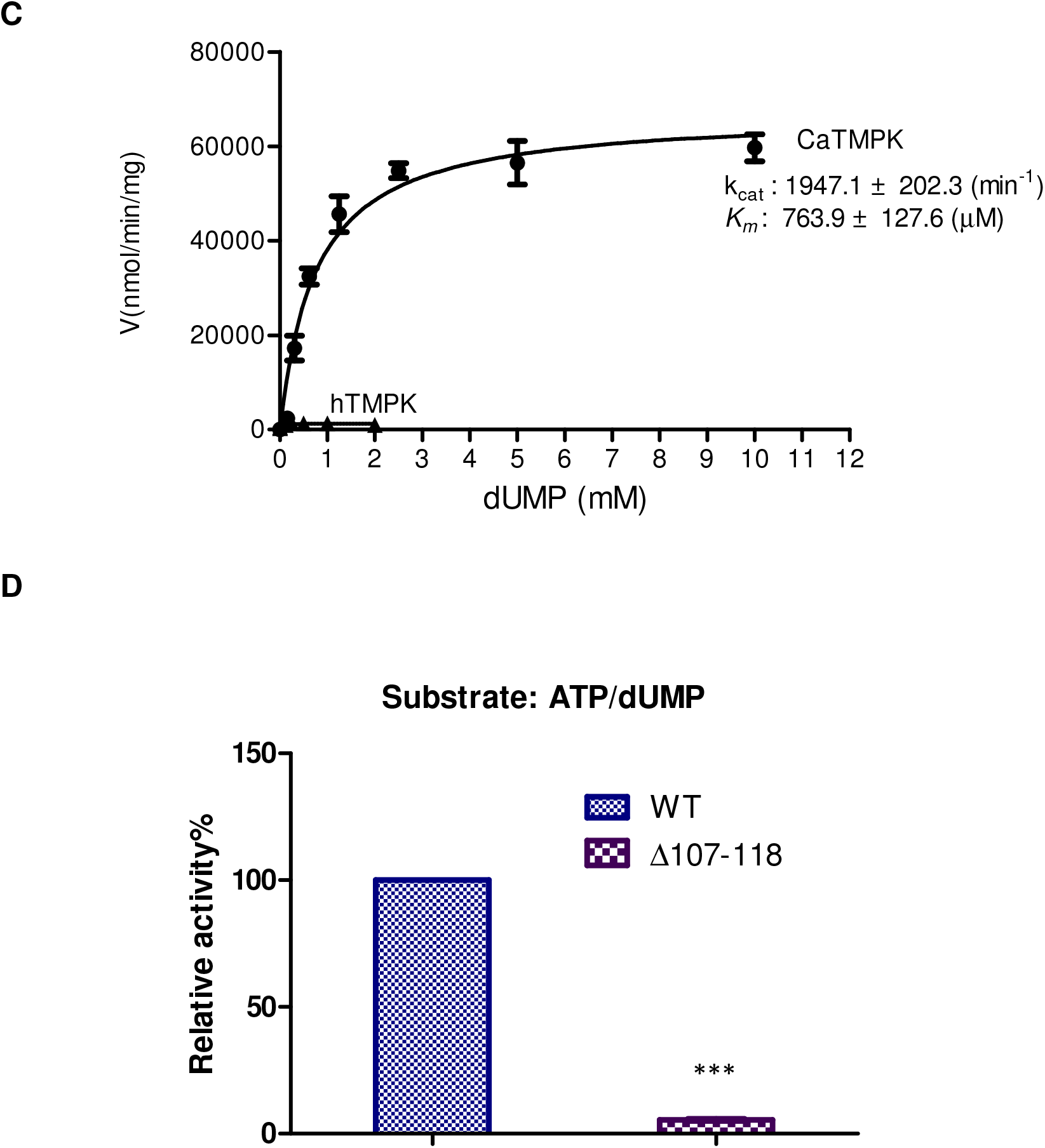
The role of the Ca-loop in the substrate specificity of CaTMPK. (A) Substrate selectivity of hTMPK and CaTMPK. Purified hTMPK and CaTMPK enzymes at the indicated amounts were subjected to an isotope-labeling-based [γ-^32^P]-ATP phosphotransfer assay as described in Materials and Methods using different substrates as indicated. Thin layer chromatography (TLC) was used to separate the reaction products of hTMPK and CaTMPK. After radiography, the position of each dNMP was visualized by 254-nm UV light and marked with a circle on the X-ray film. The ^32^P-labeled dTDP and dUDP are indicated by circles. (B) Substrate selectivity between hTMPK and CaTMPK. Thymidine (dT), deoxyuridine (dU), 5-FdU, 5-FUrd at 1 mM each was incubated with purified Herpes virus thymidine kinase (HSV-TK1, 1 μg) withγ-32P-ATP in combination with hTMPK(0.2μg) or CaTMPK (0.02μg) in each reaction, followed by TLC separation of reaction products. The position of radio-labeled NMP, Pi, dNDP and ATP on TLC sheet was revealed by autoradiography. All the indicated nucleoside was converted to nucleoside monophosphate by HSV-TK1. NMP: nucleoside monophosphate; NDP, nucleoside diphosphate. (C) Steady-state kinetics of CaTMPK and hTMPK using different dUMP concentrations in the presence of 2 mM ATP. the *K_m_* and k_cat_ were derived from the Michaelis-Menten equation: ν = Vmax[S]/ (*K_m_* + [S]). Values are presented as the mean ± S.D. from at least three independent experiments. (D) Enzymatic activity of WT and Ca-loop mutant of CaTMPK using 2 mM ATP and 10 mM dUMP as the substrates. The results are expressed relative to WT CaTMPK activity, which was set to 100%. The results represent the mean ± SD, n=3. P-values were determined by Student’s t-test (***P<0.001).

### Distinct dUTP/5-FdUTP-meditated DNA toxicity in CaTMPK- and hTMPK-expressing *S. cerevisiae*

Thymidylate synthase (TS) is the enzyme responsible for converting dUMP to dTMP. Upon uptake, 5-fluorouracil (5-FU) and 5-fluorouridine (5-FUrd) are metabolically converted to 5-FUMP and 5-FdUMP [17], the latter of which is a suicide inhibitor of TS. Therefore, treatment with 5-FU or 5-FUrd leads to dUMP accumulation. A previous study has shown that 5-FU-induced death in *S. cerevisiae* is reversed by deletion of uracil-DNA-glycosylase (*UNG1*) [18]. This finding suggests that 5-FU treatment causes misincorporation of dUTP, the removal of which by Ung1 leads to DNA toxicity and cell death. Because CaTMPK is hyperactive in the conversion of dUMP to dUDP, we then asked whether CaTMPK and hTMPK can mediate the differential toxicity of 5-FU and 5-FUrd in *S. cerevisiae*. *RWY-42-22B*, harboring a temperature-sensitive mutation of *cdc8^ts^*, is an *S. cerevisiae* strain that can grow at 23°C normally but is not viable at 30°C. The growth of this strain at nonpermissive temperatures was rescued by expression of hTMPK or CaTMPK without a significant difference in growth rate (S2 Fig.). To avoid complications associated with endogenous *cdc8^ts^*, two strains expressing hTMPK or CaTMPK were further engineered by replacement of *cdc8^ts^* with *HIS3*. We tested the sensitivity of these two strains to 5-FU. Interestingly, the growth of yeast expressing CaTMPK was obviously suppressed along a gradient increase in 5-FU concentration (0-2.5 μM) in agar plates; in contrast, only a slight response was observed in hTMPK-expressing yeast (Fig 4A). To determine whether 5-FU-induced death in the CaTMPK-expressing strain was associated with misincorporation of dUTP/5-FdUTP, *ung1* was deleted from this strain for the 5-FU sensitivity assay. The results showed that deletion of *ung1* allowed the CaTMPK strain to survive in 5-FU-containing medium (Fig 4A). Similar results were observed in 5-FUrd gradient (0-50 μM) agar plates. A high concentration of 5-FUrd was required for suppression of CaTMPK-expressing yeast growth, most likely due to the lower activity of uridine permease than uracil permease in *S. cerevisiae*. Nevertheless, *ung1* deletion abrogated the 5-FUrd response in CaTMPK-expressing yeast. In conclusion, expression of CaTMPK increases the susceptibility of yeast to 5-FU and 5-FUrd via dUTP/5-FdUTP-mediated DNA toxicity (Fig 4B).

**Fig 4.**
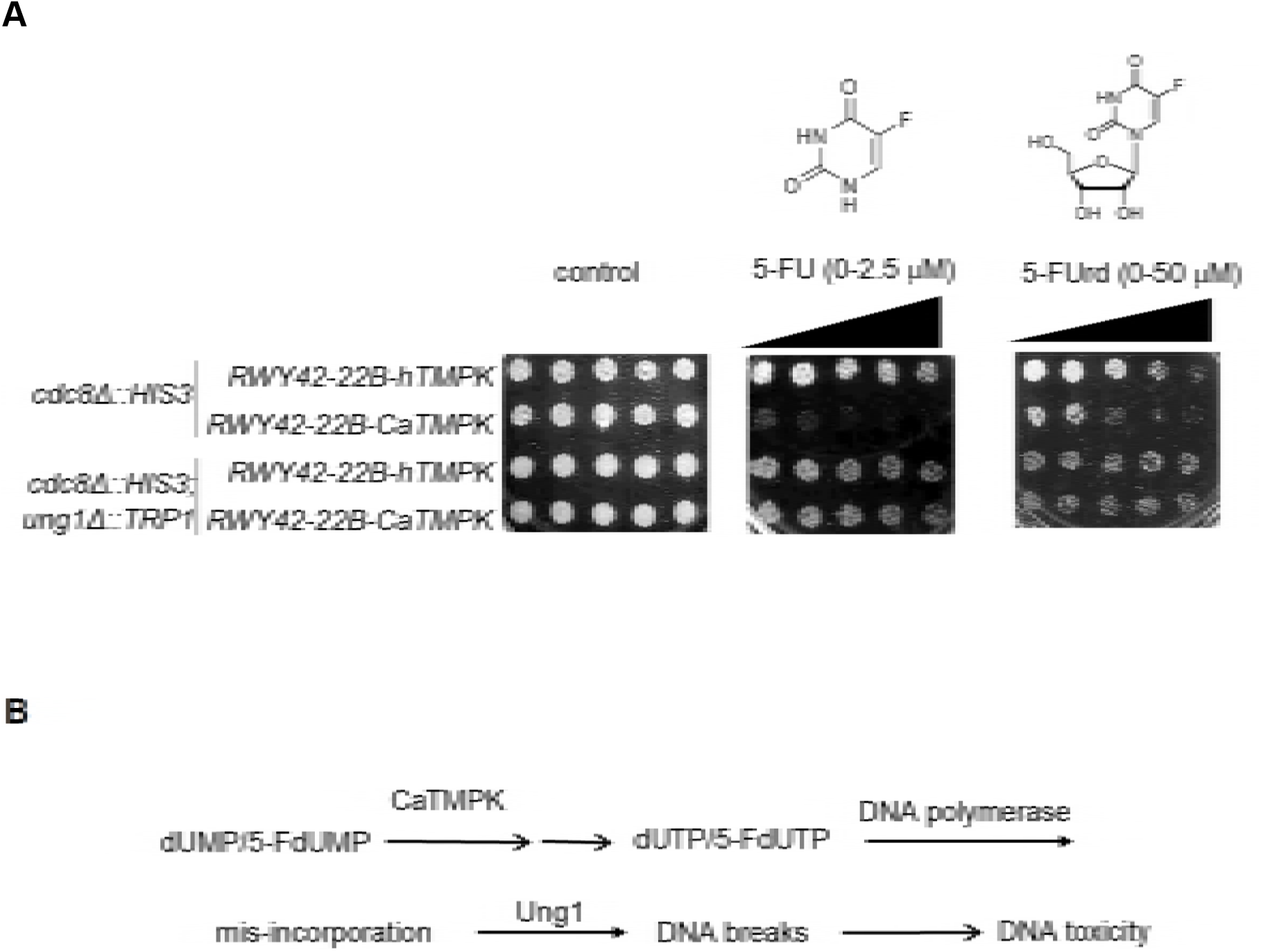
5-FU and 5-FUrd sensitivity of yeast expressing hTMPK and CaTMPK. (A) *S. cerevisiae* strains lacking endogenous *cdc8^ts^* alone or in combination with *ung1*, as indicated, and expressing hTMPK and CaTMPK were used for spot assays on agar plates in the absence or presence of an increasing gradient of 5-FU and 5-FUrd concentrations. Each spot contained 200 cells, and the plates were incubated at 30°C for 3 days. (B) The biochemical pathway responsible for 5-FU/5-FUrd-induced DNA toxicity in yeast via CaTMPK. CaTMPK mediates dUTP/5-FdUTP formation in yeast cells treated with 5-FU or 5-FUrd, leading to uracil misincorporation in DNA. Treatment with uracil-DNA glycosylase 1 (Ung1) led to the generation of excess DNA breaks, resulting in toxicity.

### Sensitivity of *C. albicans* to 5-FUrd with other drugs

5-Flucytosine (5-FC) is an antifungal pyrimidine drug used for the treatment of *Candida* infection. Upon uptake, 5-FC is converted to 5-FU by cytosine deaminase in fungi. We compared the cytotoxic effects of 5-FU, 5-FC and 5-FUrd on *C. albicans* growth. The concentration of 5-FUrd that suppressed the growth of the *HLC54* strain was in the nM range, while that of 5-FC was in the μM range. Thus, the antifungal activity of 5-FUrd is more potent than that of 5-FC. Notably, *C. albicans* growth was unaffected by 50 μM 5-FU (Fig 5A). The low 5-FU sensitivity of *C. albicans* is probably due to the poor uracil permease activity [19, 20].

**Fig 5.**
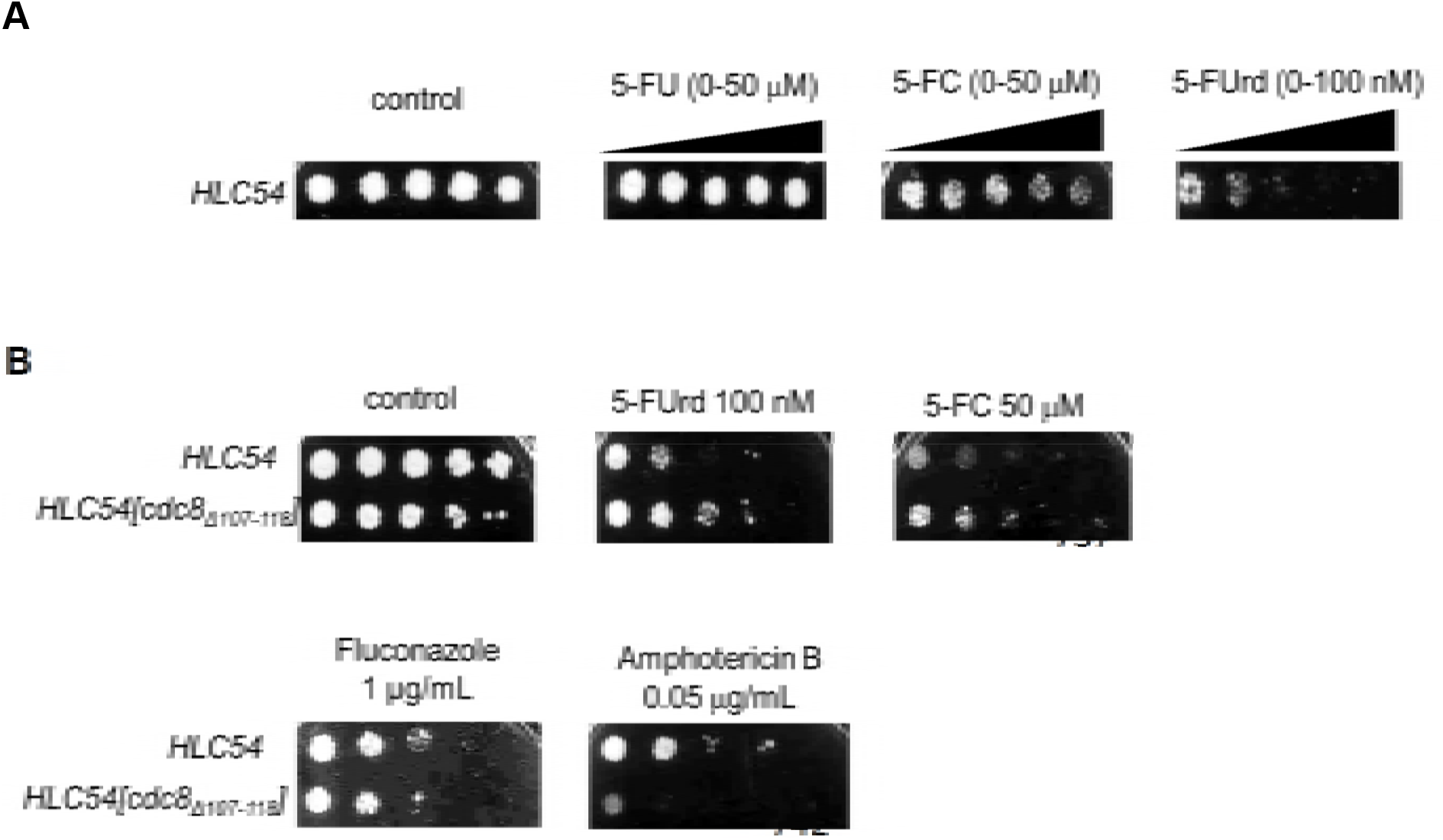
The Ca-loop affects the sensitivity of *C. albicans* to 5-FUrd and 5-FC. (A) Spot tests of the sensitivity of the *HLC54* strain to fluoropyrimidine drugs. Two hundred cells were spotted on SD-uracil plates containing increasing concentrations of 5-FU, 5-FC and 5-FUrd and incubated at 30°C for 3 days. (B) Five-fold serial dilutions of the parental *HLC54* and the *HLC54[cdc8_△107-118_*] mutant were spotted on SD-uracil plates containing the vehicle, 5-FUrd (100 nM), 5-FC (50 μM), fluconazole (1 μg/mL) or amphotericin B (0.05 μg/mL).

Given the contribution of the Ca-loop to the utilization of dUMP and 5-FdUMP as substrates of CaTMPK, we next investigated whether deletion of the Ca-loop could change the susceptibility of *C. albicans* to 5-FUrd. The 5-FUrd susceptibility was then compared in the *HLC54* and *HLC54[cdc8_Δ107-118_]* strains. Consistent with the growth rate analysis performed in liquid culture (Fig 2B), the spot assay with serial dilutions on a control plate demonstrated that the growth rate of *HLC54[cdc8_Δ107-118_]* was slower than that of *HLC54*. In contrast to the control plate, the presence of 100 nM 5-FUrd caused more pronounced growth suppression of the parental *HLC54* strain than of *HLC54[cdc8_Δ107-118_]*. Because 5-FC is intracellularly metabolized to 5-FU, a similar difference was observed between these two strains on the plate containing 50 μM 5-FC (Fig 5B). Fluconazole and amphotericin B are antifungal drugs that target ergosterol [20]. The growth responses of *HLC54* and *HLC54[cdc8_Δ107-118_]* to fluconazole were similar, whereas in the amphotericin plate, growth suppression of *HLC54[cdc8_Δ107-118_]* was more pronounced than that of the parental strain. Thus, the Ca-loop of *CDC8* makes a unique contribution fluoropyrimidine susceptibility in *C. albicans*.

### Different pathogenic *C. albicans* isolates from patients are sensitive to 5-FUrd

5-FUrd has not been considered as a drug for the treatment of *Candida* infection, probably because a high dosage of 5-FU is required. The acquisition of drug resistance in *C. albicans* has been a challenge in medical care [4, 21-24]. Given the high potency of 5-FUrd in suppressing *HLC54*, we then assessed 5-FUrd sensitivity in pathogenic *C. albicans* strains. The MIC_50_ of 5-FUrd for *ATCC90028* and *SC5314* by CLSI method [25] were 0.03 and 0.04 μg/mL (Table 2), respectively.

**Table 2.** Minimum inhibitory concentrations of 5-FUrd and antifungal drugs of clinical used for pathogenic and drug-resistant isolates of *Candida albicans^a^*.

| strain | MIC(μg/mL) | | | | | |
| --- | --- | --- | --- | --- | --- | --- |
|  | 5-FUrd<sup>b</sup> | 5-FC<sup>c</sup> | Posaconazole<sup>c</sup> | Voriconazole<sup>c</sup> | Itraconazole<sup>c</sup> | Fluconazole<sup>c</sup> |
| ATCC90028 | 0.030±0.011 | 0.5 | 0.06 | 0.015 | 0.06 | 1 |
| SC5314 | 0.044±0.003 | 0.06 | 0.015 | 0.015 | 0.06 | 0.5 |
| F2016a067 | 0.049±0.001 | 0.03 | 0.03 | 0.03 | >16(NW)<sup>d</sup> | 64(R)<sup>e</sup> |
| F2016f024 | 0.031±0.003 | 0.03 | 0.25 | 0.5 | 0.12 | 8(R) |
| F2016g048 | 0.063±0.012 | 0.03 | 1(R) | >8(R) | 1 | 256(R) |
| F2015f019 | 0.036±0.008 | 0.03 | 0.12 | 0.06 | 0.12 | 4 |
| F2014a093 | 0.021±0.001 | 0.03 | >8(R) | >8(R) | >16(NW) | >256(R) |
| F2014f090 | 0.028±0.002 | 0.12 | >8(R) | 0.25 | >16(NW) | 2 |
| F2015b076 | 0.026±0.001 | 0.03 | 1 | >8(R) | 0.5 | 256(R) |
| F2017g023 | 0.043±0.001 | >64(NW) | 0.03 | 0.004 | 0.06 | 0.5 |
| F2016c087 | 0.021±0.008 | >64(NW) | 0.06 | 0.015 | 0.12 | 0.5 |
| F2016e085 | 0.076±0.020 | 8(NW) | 0.015 | 0.004 | 0.03 | 0.06 |
| F2017c022 | 0.018±0.002 | >64(NW) | 0.015 | 0.004 | 0.03 | 0.25 |
| F2017c071 | 0.027±0.003 | 4(NW) | 0.015 | 0.004 | 0.03 | 0.25 |

**Abbreviation:** 5-FC, 5-flucytosine; (R), Resistance; (NW), Non-wild-type strain
<sup>a</sup>*Candida* human blood isolates were collected from patients with bloodstream infection due to *Candida albicans* during a prospective observational study which was approved by the Research Ethics Committees (NTUH-201103121RB, NTUH-201502034RIND).
<sup>b</sup>Minimum inhibitory concentrations (MICs) of 5-FUrd were measured by broth microdilution method as described in materials and methods and determined in accordance with the guidelines in Clinical and Laboratory Standards Institute (CLSI) document M27[25].
<sup>c</sup>MICs were determined by the microdilution colorimetric Sensititre YeastOne SYO-09 panel (TREK Diagnostic Systems, Cleveland, OH, USA) as described in materials and

We further tested the 5-FUrd sensitivity of a number of pathogenic *C. albicans* isolates from patients. The MIC_50_ of 5-FUrd was 0.03 μg/mL for the drug-susceptible isolate and was 0.01-0.08 μg/mL for five 5-FC-resistant isolates, designated non-wild-type strains. For azole-resistant isolates, 5-FUrd continued to efficiently inhibit growth in the range 0.02-0.07 μg/mL. Because 5-FC is intracellularly metabolized to 5-FU, we asked why the 5-FC-resistant isolates remained sensitive to 5-FUrd. Several mutations in 5-FC resistant strains were identified in the enzymes involved in the formation of 5-FUMP from 5-FC, including purine-cytosine permease (Fcy2), cytosine deaminase (Fca1) and uracil phosphoribosyltransferase (Fur1) [23]. These 5-FC-resistant but 5-FUrd-sensitive strains might have mutations in these genes. 5-FUrd is transported into *C. albicans* through uridine permease (Fui1) and is then phosphorylated to 5-FUMP by uridine kinase (Urk1) [26]. Presumably, activation of 5-FUrd metabolism by the Fui1 and Urk1 pathways can overcome the defects in genes mediating 5-FC activation, namely, Fcy2, Fca1 and Fur1, leading to toxicity, as depicted in Fig 6. One important effect of antifungal drugs is host liver toxicity. We then evaluated the potential toxicity of 5-FUrd in human HepG2 cells and found that the IC50 of 5-FUrd was 4 μg/mL (S3 Fig.). Taken together, these results suggest that 5-FUrd might be a suitable option for antifungal treatment.

**Fig 6.**
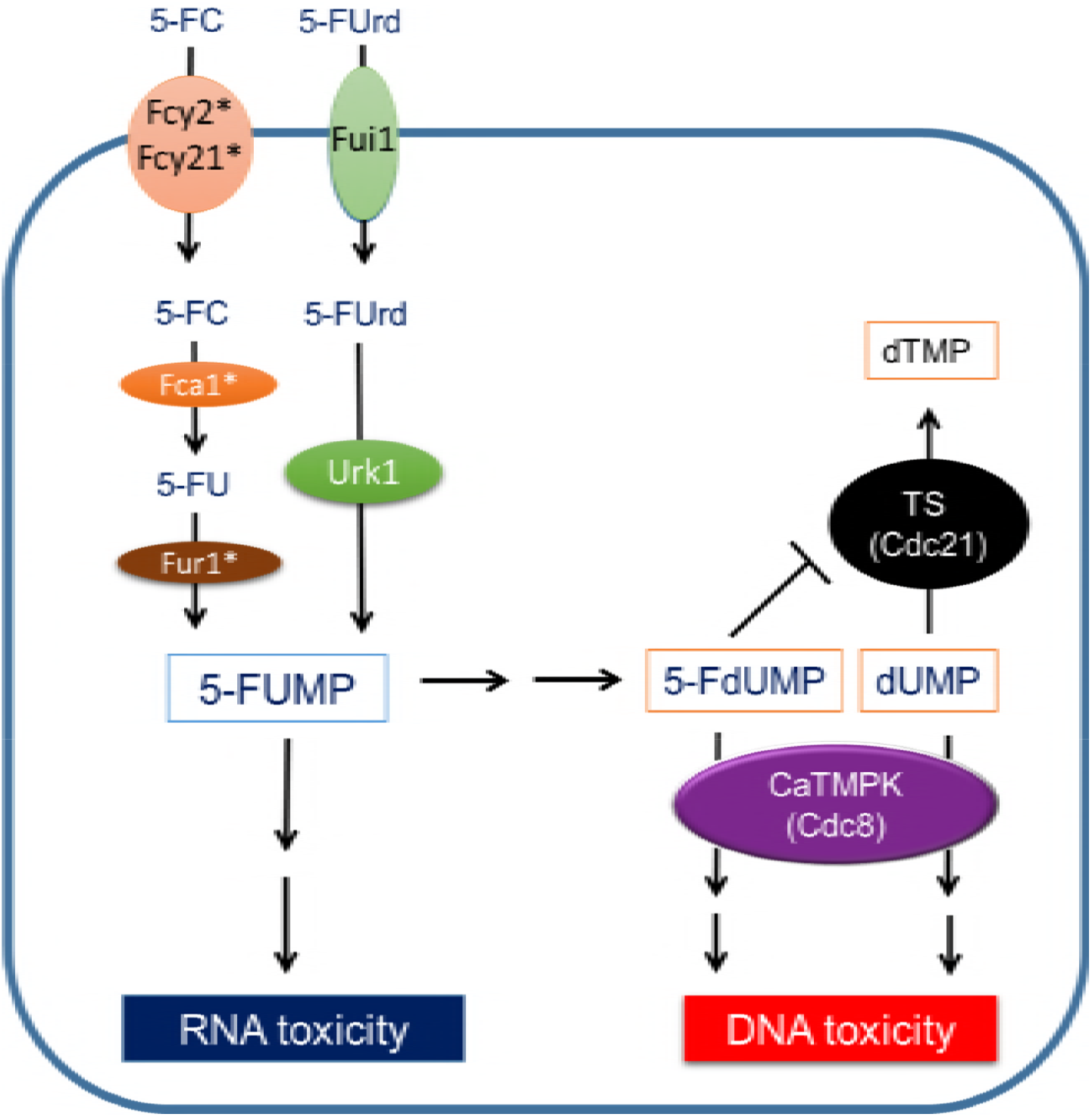
Mechanism underlying the overcoming of 5-FC resistance by 5-FUrd in *C. albicans*. Differences in the 5-FC and 5-FUrd activation pathways and toxic effects of CaTMPK in *C. albicans*. Nomenclature of the enzymes and coding genes in *C. albicans*: Fca1, cytosine deaminase; Fcy2, purine-cytosine permease; Fcy21, pseudogene paralog of Fcy2; Fur1, uracil phosphoribosyltransferase; Cdc21, thymidylate synthase; Cdc8: thymidylate kinase (CaTMPK). Asterisks indicate genes that have been found to be mutated in 5-FC resistance.

## Discussion

Members of the TMPK enzyme family have been categorized into type I and type II enzymes. Both hTMPK and CaTMPK are type I enzymes [7]. The in vitro analyses of this study highlight two major differences between of the purified hTMPK and CaTMPK. First, the catalytic efficiency of CaTMPK is 15–fold higher than hTMPK. Second, CaTMPK is highly active in the conversion of dUMP to dUDP and 5-FdUMP to 5-FdUDP. Consistent with the data from biochemical analyses, we provide evidence that this Ca-loop-mediated hyperactivity determines the growth rate of *C. albicans*, suggesting the potential of targeting this site to control infection of this pathogen. Furthermore, the differences in the capacity to use dUMP and 5-FdUMP as substrates support the finding that CaTMPK, but not hTMPK, can mediate dUTP/5-FdUTP toxicity under 5-FU and 5-FUrd treatment in *S. cerevisiae*. Although 5-FU is not useful for the treatment of *C. albicans* infection probably due to the lack of transporter, we observed that nonpathogenic and pathogenic strains of *C. albicans* are very sensitive to 5-FUrd. We propose that the utilization of dUMP and 5-FdUMP as substrates by CaTMPK increases the 5-FUrd susceptibility of *C. albicans*. In addition, our finding also open a new question whether the high activity of CaTMPK in dUMP utilization might participate in the mechanism of stress-induced genome evolution [28, 29] of *C. albicans* by uracil misincorporation in DNA, thereby facilitating the development of drug resistance.

CRISPR-mediated deletion was performed to generate a *C. albicans* strain lacking the Ca-loop of CaTMPK led to markedly reduced growth. This result indicated that the hyperactivity of CaTMPK in dTDP synthesis is an important factor affecting the growth rate of *C. albicans*. Notably, in this study, we also used the *S. cerevisiae* expression system to evaluate differences between the effects of hTMPK and CaTMPK on growth and drug sensitivity. Despite the considerable differences in catalytic rates, the growth rates of *S*. *cerevisiae* expressing CaTMPK and hTMPK were very similar. This observation is consistent with a previous report that demonstrated that decreased thymidylate kinase activity did not affect the growth of *S. cerevisiae* [30]. Apparently, *C. albicans* and *S. cerevisiae* have different requirements for TMPK activity for growth maintenance. Notably, we observed a striking difference in 5-FU and 5-FUrd toxicity between *S. cerevisiae* strains expressing hTMPK or CaTMPK. Intracellular 5-FU is converted to 5-FdUMP, which is an inhibitor of TS, thereby blocking the synthesis of dTMP from dUMP, which in turn results in the accumulation of dUMP. Because dUMP is not a good substrate for hTMPK, cells expressing hTMPK would have less dUTP accumulation. In contrast, dUMP and 5-FdUMP are good substrates for CaTMPK, and dUDP formation by this enzyme might facilitate dUTP accumulation, to leading to misincorporation in DNA. Uracil is removed from DNA by the action of Ung1 [18], thereby generating DNA breaks. Excess uracil incorporation therefore leads to DNA toxicity. Given that deletion of *ung1* abolished 5-FU and 5-FUrd toxicity in *S. cerevisiae* expressing CaTMPK, it is very clear that high activity of CaTMPK in the utilization of dUMP and 5-FdUMP is responsible for DNA toxicity of 5-FU and 5-FUrd.

In this study, our data revealed 5-FUrd to be a potent drug for the treatment of *C. albicans* infection. This proof-of-concept study showed that a number of *C. albicans* isolates that were resistant to azoles or 5-FC were susceptible to 5-FUrd at a dose that had little toxicity in human hepatoma cells. We proposed that 5-FUrd could overcome 5-FC resistance in these strains because the metabolic activation pathways of 5-FUrd and 5-FC are different. Therefore, 5-FC resistance resulting from mutations in genes mediating the metabolic activation of 5-FC might not affect 5-FUrd toxicity. In summary, this report reveals the biochemical differences between the essential enzyme TMPK from *C. albicans* and host (human) TMPK due to the Ca-loop, highlighting this enzyme to be a new drug target site. Moreover, we found that 5-FUrd is indeed a potent inhibitor of *C. albicans* growth and that the Ca-loop is involved in 5-FUrd toxicity. Notably, 5-FUrd is able to overcome 5-FC and multidrug resistance in pathogenic *C. albicans* isolates.

## Methods

### Media and chemicals

All *S. cerevisiae* strains in this study were grown in synthetic defined (SD) medium containing 2% glucose, 6.7 g/L yeast nitrogen base without amino acids and 0.77 g dropout (DO) supplements –Ura (Clontech). *C. albicans* were grown in YPD (1% yeast extract, 2% bactopeptone, and 2% glucose) or SD medium at 30 °C. For drug susceptibility analysis, *C. albicans* were grown in RPMI1640 (Sigma Aldrich), which buffering with MOPS (3-(N-Morpholino) propanesulfonic acid, 4-Morpholinepropanesulfonic acid; Sigma Aldrich). For solid media, 2% agar was added. [γ-^32^P] ATP (10 μCi/μL) was from Moravek Biochemicals. Polyethyleneimine (PEI) cellulose F plastic sheets (layer thickness 0.1 mm) for TLC was from Millipore. Yeast nitrogen base, thymidine, deoxyuridine, dAMP, dCMP, dGMP, dTMP, dUMP, 5-Fluorouridine, 5-Fluorouracil, 5-Fluorocytocine, NADH, lysozyme, pyruvate kinase and Lactate dehydrogenase were from Sigma Aldrich. Nourseothricin (Nat) was purchase from Jena Bioscience.

### Plasmids construction

*C. albicans CDC8* in pUC57 was purchased from GenScript (#5001191-1). Due to the difference in CTG codon usage for serine in *C. albicans* and leucine in *E. coli* and *S. cerevisiae*, the expression of *CDC8* gene in *E. coli* and *S. cerevisiae* would cause S68L mutation. This codon of *CDC8* gene was mutated to TCG by site-directed mutagenesis to maintain the correct protein sequence termed CaTMPK when expressed in *E. coli* and *S. cerevisiae*. CaTMPK and hTMPK cloned to pGEX-2T were used to produce recombinant protein CaTMPK and hTMPK in *E. coli*. CaTMPK deleted of 107-118 amino-acid were generated by site-directed PCR mutagenesis in pGEX-2T-CaTMPK. For expression in *S. cerevisiae*, PCR-amplified hTMPK and CaTMPK fragments were each cloned to *pRS416* plasmid (a gift from M.Y Chen, National Yang-Ming University, Taiwan) containing the *TEF1* promoter and *CYC1* terminator. The plasmid pV1025 and pV1090 (kindly provided by Valmik K. Vyas, Cambridge) were used for CRISPR genome editing in *C. albicans*. The plasmid pV1090-17 carrying sgRNA sequence targeting *CDC8* in *C. albicans* was cloned at BsmBI restriction sites of the plasmid pV1090, and the plasmid pV1025 contains Cas9 for expression in *C. albicans*. Plasmids and primers in this study are listed in Table S3 and Table S4.

### Establishment of *S. cerevisiae* and *C. albicans* strains

*S. cerevisiae* strains *RWY42-22A* (Mata *CDC8*) and *RWY42-22B* (Matα *cdc8-1*) are provided by Raymund J. Wellinger (Université de Sherbrooke). *RWY42-22B* (Matα *cdc8-1*) contains temperature-sensitive (ts) mutation allele of *cdc8 (cdc8-1)*, which is able to grow at 23°C but not at 30°C. The isogenic *CDC8* strain (WT), RWY42-22A, is able to grow at both temperatures. *RWY42-22B* was further transformed with plasmid *pRS416-TEF1-hTMPK-CYC1* and *pRS416-TEF1-CaTMPK-CYC1*, respectively, and replaced endogenous *cdc8-1* with *HIS3* gene, thus generating *RWY42-22B-1A* and *RWY-42-22B-1B*. The endogenous *UNG1* gene of *RWY42-22B-1A* and *RWY42-22-1B* was replaced by *TRP1* marker, which was amplified from the plasmid *pRS314* (provided by M.Y Chen, National Yang-Ming University, Taiwan), thus generating *RWY42-22B-2A* and *RWY42-22B-2B* strains.

Non-pathogenic *C. albicans* strain, *HLC54* [31] (provided by Hsiu-Jung Lo, National Health Research Institutes, Taiwan), was used to generate endogenous Δ107-118 mutation in endogenous *CDC8* locus by a CRISPR system [16]. Briefly, pV1090-17 and pV1025 (20 μg) were linearized before transformation by a modified LiAC method[32]. For Ca-loop deletion in endogenous *CDC8* locus, donor template (378bp) were amplified from the plasmid pGEX-2T-CaTMPKΔ107-118 and purified for transformation along with linearized pV1090-17 and pV1025. The transformants were spread on YPD plate containing Nat at 250 μg/ml and incubated at 30°C for 3 days. The colonies grown on Nat plate were isolated for genomic DNA extraction followed by PCR examination. All strains in this study are listed in Table S2.

### Purification of recombinant proteins

pGEX-5X-HSV-TK1, pGEX-2T-hTMPK and pGEX-2T-CaTMPK (WT/Ca-loop mutants) were each transformed into *E. coli* BL21 strain. A single clone was inoculated in 15 ml of LB broth and cultured overnight at 37°C. 10 mL of the overnight culture was dilute to 1000 mL and cell growth was continued at 37°C until OD_600_ reached 0.5. To induce hTMPK and CaTMPK expression, IPTG (isopropyl β-D-1-thiogalactopyranoside) was added to a final concentration of 0.5 mM and the culture was incubated for another 3.5 hours at 37°C and 32°C, respectively. For HSV-TK1 induction, 0.5 mM IPTG was added, and culture was incubated at 27°C for 16 hr. Cells were harvested by centrifugation and re-suspended in 20 mL of lysis buffer containing 1% NP40, 50 mM Tris-HCl pH7.5, 150 mM NaCl, 5 mM MgCl_2_, 1 mM DTT, 1mM PMSF and 1 mM protease cocktail before being disrupted by sonication. Following centrifugation, 2 mL of glutathione Sepharose beads was added to the clarified cell lysate and incubated with gentle shaking at 4°C for 1.5 hours. Beads were washed five times with a buffer containing 0.5% NP40, 50 mM Tris-HCl pH7.5, 150 mM NaCl, 5 mM EDTA, 1 mM DTT, 1mM PMSF and 1 mM protease cocktail. The GST-tag was then cleaved from the target proteins by thrombin.

### *In vitro activity* assay (NADH-coupled assay)

Activity assay was measured using spectrophotometric method by coupling ADP formation to the oxidation of NADH catalyzed by pyruvate kinase and lactate dehydrogenase. Reaction mixture for each set in 100μL was contained 500 μM ATP, 500 μM dTMP (for measurement of dUMP activity, 2 mM ATP and 10 mM dUMP),100 mM Tris-HCl, 100 mM KCl and 10 mM MgCl_2_, 500 μM Phosphoenolpyruvate, 250μM NADH, 4 unit of pyruvate kinase and 5 unit of lactate dehydrogenase. Reaction was initiated by adding purified proteins (0.4 μg hTMPK or 0.015 μg CaTMPK) and was measured the reduction of NADH by 340 nm. For hTMPK inhibitor assay, both of hTMPK (0.4 μg) and CaTMPK (0.015 μg) were preincubated with compounds at indicated concentrations for 10 min, followed by adding reaction mixtures as described above and measuring OD_340_. The *K_m_* for dTMP and dUMP were measured by various concentrations of substrates using 500 μM ATP (2 mM for dUMP). This concentration was followed previous work [15]. The *K_m_* for ATP was measured by varying concentrations using 250 μM dTMP. The *K_m_* values were obtained by fitting Michaels–Menten equation.

### ^32^P- phosphate transfer assay

The substrate selectivity of hTMPK and CaTMPK were measured by ^32^P-Phosphate transfer assay. TMPK (0.2 μg) and CaTMPK (0.02μg) after incubation with dNMP at the 1mM for 10 min were added to the TMPK reaction mixture containing 0.05 μM [γ-^32^P] ATP (10 μCi/μL), 50 μM ATP, 100 mM Tris-HCl, 100 mM KCl and 10 mM MgCl_2_. The assay was terminated by heating at 95°C. 2μl of reaction mixture was spotted onto PEI-cellulose thin-layer chromatography (TLC). After air dry, 2 μl the corresponding dNMP (20 mM) also spotted on TLC for separation by 2 M acetic acid/ 0.5M LiCl. The position of nucleotides on TLC sheet was visualized by UV, followed by autoradiography for assessing ^32^P-Phosphate transfer.

### Yeast growth curve and drug susceptibility tests

Cells at 10^4^ cells/ mL in SD-Ura medium were plated onto a clear flat-bottom 96-well plate and incubate at 30°C in a TECAN SPARK plate reader to obtain growth curve by OD_600_ reading every 30 minutes.

Minimum inhibitory concentrations (MICs) of 5-FUrd were determined by broth microdilution method in accordance with the guidelines in Clinical and Laboratory Standards Institute (CLSI) document M27[25], using RPMI 1640 medium containing 0.165 M MOPs (pH 7.0) and 0.2% glucose, an inoculum of 10^3^ cells/well, and incubated for 24 hr at 35°C, followed by OD_600_ reading by TECAN SPARK plate reader. The MIC_50_ of 5-FUrd was determined as the lowest dosage of 5-FUrd causing 50% inhibition of the growth by using the concentrations in the range of 0.004-2μg/mL. MICs of 9 antifungal agents of human use (5-FC; four azoles: posaconazole, voriconazole, itraconazole, fluconazole; and three echinocandins: anidulafungin, caspofungin and micafungin) were determined by the microdilution colorimetric Sensititre YeastOne SYO-09 panel (TREK Diagnostic Systems, Cleveland, OH, USA); in accordance with the manufacturer’s instructions. MICs values were determined visually, after 24 hours of incubation, as the lowest concentration of drug that caused complete inhibition (amphotericin B) or a significant diminution (≥50% inhibition; flucytosine, azoles and echinocandins) of growth relative to that of the growth control.

For drug susceptibility assay on solid medium plate, *S. cerevisiae* or *C. albicans* colonies were inoculated in 5 ml of SD selection mediums, and grown overnight to logarithmic phase at 30°C. For gradient plate, cells were counted and 200 cells were spotted onto solid medium containing an increasing concentrations of drugs in the agar. For plates with fixed concentrations of drug, the cultures were adjusted to 0.1 by OD_600_, then serial diluted with five-fold before spotted on plates. The plates were incubated at 30°C for 2–3 days.

### Statistical analysis

Data are presented as the mean ± standard error of the mean. Statistical comparison of means was performed using a two-tailed unpaired Student’s t-test.

## Acknowledgments

We thank Hsueh-Tzu Shih for performing 5-FUrd cytotoxicity assay in HepG2 and LiFan Chen for technical assistance of CLSI tests. We thank Valmik K. Vyas (Massachusetts Institute of Technology, Cambridge) for kindly providing *C. albicans* CRISPR system and protocol. We thank Mei-Yu Chen (National Yang-Ming University, Taipei, Taiwan) for providing expression vectors in *Saccharomyces cerevisiae*.

## Supporting information

**S1 Fig. Deletion of Ca-loop at gene *CDC8* locus in *HLC54* strain by CRISPR**. (A) the CRISPR system used in this study. (B) The sgRNA targeting region designed at Ca-loop sequence (purple), the repaired template contained the middle region of *CDC8* gene without Ca-loop sequence. Linearized DNA fragments of Cas9, sgRNA and donor template were co-transformed into *HLC54* and selected by Nourseothricin (Nat). (C) The genomic DNA was extracted from the survived clones on Nat^R^ plate for PCR reaction. The PCR product gave 439bp from wild-type *CDC8*, while 403bp was from the clone with 107-118 deletion. Forward primer: ATCAAGCAGCTCATTTGTTATTTCTGGC; Reverse primer: GATTTCCCATAGTTGTGTAGTTACTTGATC. (D) Sanger sequencing of the region spanning 107-118 of *cdc8* in *HLC54[cdc8_Δ107-118_]* clone.

**S2 Fig. Growth of yeasts at 23°C and 30°C**. 10-fold serial dilution of yeast cells on SD-Uracil plates, which were incubated at 23°C and 30°C for 3 days. *RWY-42-22A* contains WT *CDC8*, whereas *RWY-42-22B* contains mutant form *cdc8*.

**S3 Fig. Effect of 5-FUrd on HepG2 cell viability assay**. Cells seeded on 96-well plates for 24 hr were treated with various concentrations of 5-FUrd. After 72 hr, cells were measured by CellTiter-Glo (Promega). Results normalized with no drug, and set to 100%. Result represent in means ± SD, n=4.

**S1 Table. Yeast strains in this study**

**S2 Table. Plasmids in this study**

**S3 Table. Oligonucleotides in this study**

